# SARS-CoV-2 Omicron variant is more stable than the ancestral strain on various surfaces

**DOI:** 10.1101/2022.03.09.483703

**Authors:** Alex Wing Hong Chin, Alison Man Yuk Lai, Malik Peiris, Leo Lit Man Poon

## Abstract

The Omicron BA.1 SARS-CoV-2 variant of concern spreads quickly around the world and outcompetes other circulating strains. We examined the stability of this SARS-CoV-2 variant on various surfaces and revealed that the Omicron variant is more stable than its ancestral strain on smooth and porous surfaces.

---

The newly emerged Omicron SARS-CoV-2 variant of concern (VOC) is highly transmissible in humans. It outcompetes other previously known variants and dominates in different geographical locations in recent months (1). Its spike protein has more than 30 mutations compared to the ancestral strain (2). A recent structural study indicates its spike protein is more stable than the ancestral strain (3). This prompts us to hypothesize that Omicron VOC is also more stable on different surfaces. We previously showed that the ancestral SARS-CoV-2 can still be infectious for several days and hours at room temperature on smooth and porous surfaces, respectively (4). Here, we report that Omicron VOC is more stable than the ancestral strain on these surfaces.

Previously described ancestral SARS-CoV-2 (PANGO lineage A) and Omicron VOC (PANGO lineage BA.1) were used in this study (5, 6). Their stability on different surfaces were tested using our previously described protocol by us (4, 7). In brief, a 5 μl droplet of each virus (10^7 TCID50/ml) was applied on different surfaces with a dimension of 1×1 cm^2^. The treated surfaces were incubated at room temperature (21-22°C) for different time points as indicated and were then immersed in viral transport medium for 30 min to recover residual infectious virus. The recovered virus was titrated by TCID50 assays using Vero-E6 as described (4, 7).

When compared to the ancestral SARS-CoV-2, the Omicron BA.1 variant was shown to be more stable on all studied surfaces (Table). At 2 days post-incubation, the infectious viral titres of ancestral SARS-CoV-2 recovered from stainless steel, polypropylene sheet and glass reduced by 99.91%, >99.86% and 99.9%, respectively. No infectious ancestral SARS-CoV-2, except in one out of three treated glass samples, could be recovered on day 4 post-incubation. In contrast, the Omicron variant could still be recovered from these treated surfaces on day 7 post-incubation. Infectious Omicron variant virus recovered from treated stainless steel, polypropylene sheet and glass on day 7 post-incubation reduced by 98.19%, 99.65% and 98.83%, respectively. Thus, the infectious titres of the Omicron variant were not reduced by 3 log10 units on any of these smooth surfaces at our last study time point.

**Table:** Stability of the ancestral SARS-CoV-2 and Omicron variant on different surfaces.

**114 Table: Stability of the ancestral SARS-CoV-2 and Omicron variant on different surfaces**
| Materials | Time of incubation | Ancestral SARS-CoV-2 |  | Omicron variant |  |
| --- | --- | --- | --- | --- | --- |
|  |  | Mean<br>Log<sub>10</sub>(TCID<sub>50</sub>/ml)<br>± S.D. | %<br>Reduction<br>in viral<br>titre | Mean<br>Log<sub>10</sub>(TCID<sub>50</sub>/ml)<br>± S.D. | %<br>Reduction<br>in viral<br>titre |
| Stainless steel | 0 | 5.02 ± 0.39 | NA | 5.35 ± 0.18 | NA |
|  | 3 h | 4.21 ± 0.36 | 85.15% | 4.82 ± 0.23 | 69.78% |
|  | 6 h | 3.73 ± 0.10 | 95.80% | 4.62 ± 0.31 | 79.86% |
|  | 1 day | 2.99 ± 0.17 | 99.21% | 4.65 ± 0.17 | 80.28% |
|  | 2 days | 2.08 ± 0.11 | 99.91% | 4.51 ± 0.15 | 85.82% |
|  | 4 days | < | >99.93% | 3.72 ± 0.12 | 97.72% |
|  | 7 days | < | >99.93% | 3.58 ± 0.30 | 98.19% |
| Poly-propylene | 0 | 4.85 ± 0.23 | NA | 5.43 ± 0.16 | NA |
|  | 3 h | 4.12 ± 0.19 | 81.72% | 4.65 ± 0.34 | 81.27% |
|  | 6 h | 3.53 ± 0.15 | 95.43% | 4.33 ± 0.14 | 92.34% |
|  | 1 day | 3.13 ± 0.34 | 97.86% | 4.45 ± 0.23 | 89.25% |
|  | 2 days | *2.01 ± 0.10 | >99.86% | 4.34 ± 0.25 | 91.53% |
|  | 4 days | < | >99.88% | 3.97 ± 0.19 | 96.48% |
|  | 7 days | < | >99.88% | 2.95 ± 0.27 | 99.65% |
| Glass | 0 | 5.10 ± 0.24 | NA | 5.65 ± 0.28 | NA |
|  | 3 h | 4.26 ± 0.05 | 86.79% | 4.90 ± 0.15 | 83.62% |
|  | 6 h | 3.69 ± 0.11 | 96.42% | 4.52 ± 0.13 | 93.20% |
|  | 1 day | 2.83 ± 0.13 | 99.49% | 4.20 ± 0.01 | 96.84% |
|  | 2 days | 2.14 ± 0.13 | 99.90% | 4.43 ± 0.29 | 93.87% |
|  | 4 days | *1.96 ± 0.00 | >99.93% | 4.06 ± 0.16 | 97.64% |
|  | 7 days | < | >99.93% | 3.76 ± 0.10 | 98.83% |
| Tissue paper | 0 | 4.70 ± 0.22 | NA | 5.21 ± 0.14 | NA |
|  | 5 min | 3.85 ± 0.28 | 84.98% | 4.64 ± 0.70 | 53.94% |
|  | 15 min | 2.12 ± 0.14 | 99.75% | 3.72 ± 1.22 | 72.99% |
|  | 30 min | < | >99.84% | 2.92 ± 0.40 | 99.34% |
|  | 60 min | < | >99.84% | < | >99.95% |
| Printing paper | 0 | 5.21 ± 0.00 | NA | 5.34 ± 0.13 | NA |
|  | 5 min | 2.69 ± 0.16 | 99.68% | 3.26 ± 0.42 | 98.91% |
|  | 15 min | < | >99.94% | *2.20 ± 0.33 | >99.91% |
|  | 30 min | < | >99.94% | *2.16 ± 0.36 | >99.92% |
|  | 60 min | < | >99.94% | < | >99.96% |

< All the triplicates were below detection limit of the TCID<sub>50</sub> assay.
\*One or two out of three replicates were below detection limit of the TCID<sub>50</sub> assay.

The stability of the Omicron variant was also higher than ancestral SARS-CoV-2 on porous surfaces such as facial tissue paper and printing paper. On tissue paper, viable ancestral SARS-CoV-2 was no longer recoverable in 30 min after incubation. However, for the Omicron variant, viable virus could still be detected after a 30-minute incubation and the reduction in titre was less than 3 log10 units (99.34%). On printing paper, the virus titre of ancestral SARS-CoV-2 reduced by 99.68% in 5 minutes and no infectious virus could be detected after a 15-minute incubation. In contrast, the Omicron variant was more stable than the ancestral SARS-CoV-2, with viable Omicron variant virus still recovered from two out of three replicates after a 30-minute incubation.

Overall, the Omicron variant is more stable on different surfaces and materials than the ancestral strain. More evidence is needed to account for the increased transmissibility of Omicron variant observed in various community studies. The extra virus stability on surfaces may be one possible factor and should be taken into consideration when recommending control measures against the infection. A recent study revealed that an infectious dose as low as 10 TCID50 units could infect more than 50% of human subjects (8). Our findings imply that Omicron variant has an increased likelihood to be transmitted by the fomite route. Hand hygiene and frequent disinfection of common touch surfaces in public areas are highly recommended. Guidelines for disinfecting a contaminated site might also need to be reviewed (https://www.cdc.gov/coronavirus/2019-ncov/community/disinfecting-building-facility.html). One may also speculate that the enhanced stability deduced from structural studies and now demonstrated on different surfaces may be relevant for droplet or aerosol transmission of SARS-CoV-2. Interestingly, stability of avian influenza A H5N1 viruses has been shown to have an association with transmissibility of avian influenza virus between mammals by the airborne route although the mechanisms underlying this association have not been fully understood (9). Further studies on the stability of Omicron variant in droplets and aerosols are warranted.

This study has some limitations. The experiments were carried out in a laboratory with well-controlled environment. Variation in the environmental conditions would affect the rate for viral inactivation. Therefore, the time required for virus inactivation demonstrated in this study may not completely reflect all scenarios in daily life. It should also be noted that the components of the medium of the viral droplets applied in this study were different from the respiratory droplets. This could also affect the stability of the virus. Irrespective of these effects, our findings demonstrate that the Omicron variant is more stable than the ancestral SARS-CoV-2 on different surfaces and we suggest that our findings may be relevant for public health.

## Acknowledgements

This work was supported by the Health and Medical Research Fund (COVID190116) and RGC theme-based research schemes (T11-705/21-N) and InnoHK grants for C2I.

## Biographical sketch of the first author

Dr. Alex W.H. Chin is currently a Research Assistant Professor of The University of Hong Kong. His research interests include basic virology and pathogenesis of emerging respiratory viruses.

